# Comparative metagenomic profiling of seed-borne microbiomes in a landrace and a hybrid maize variety

**DOI:** 10.1101/2024.04.04.588073

**Authors:** Sarah Henaut-Jacobs, Beatriz Elisa Barcelos Cyríaco, Francisnei Pedrosa-Silva, Fabio Lopes Olivares, Thiago Motta Venancio

**Author notes:** Corresponding authors at Av. Alberto Lamego 2000, P5 sala 217; Parque California, Campos dos Goytacazes, RJ, Brazil., E-mail addresses.

## Abstract

The plant seed-borne microbiome comprises microorganisms vertically inherited from the mother plant. This microbiome is often linked to early-life protection and seedling growth promotion. Here, we compare the seed-borne bacteriomes of a commercial hybrid and a landrace maize variety. The landrace variety displays a more diverse seed-borne microbiome, featuring a variety of taxa across samples. In contrast, the microbiome of the hybrid variety is less diverse and more uniform across samples. Although both microbiomes lack a functional nitrogen fixation apparatus, we found a remarkably distinct presence of genes associated with phytohormone production and phosphate solubilization, particularly in the landrace variety. In addition, we recovered 18 metagenome-assembled genomes (MAGs), including four from potentially novel species. Collectively, our results allow a better understanding of the contrasting diversity between maize varieties and open important perspectives for designing synthetic microbial communities for agroecosystems.

## Introduction

When examining the intricate world of microscopic ecosystems, plants are dynamic habitats in their own right. Plants rely heavily on the microbial communities within and around them ^1^, engaging in myriad complex interactions that can be beneficial, antagonistic, or neutral ^2^. The plant bacteriome comprises bacteria acquired from the surrounding soil and those vertically transmitted during seed development. These vertically inherited bacteria comprise the resident or seed-borne microbiome. The seed-borne microbiome is often associated with seed-to-seedling growth transition and protection ^3^.

Factors such as plant genotype and environmental conditions can influence the plant microbiome composition and functions, emphasizing the dynamic nature of these bacterial communities ^4^. Different plant genotypes are likely to diverge in their seed-borne microbiomes due to artificial selection and recruitment of other bacteria from the soil ^5^. In addition, neighboring plants ^6^ and long-term processes such as global warming ^7^ can influence a plant microbiome. Maize is a suitable model for studying seed-borne microbiomes. Besides its great economic relevance as the second most produced agricultural commodity in the world ^8^, maize has a well-documented evolutionary and domestication history, valuable germplasm collections, and other genetic resources ^9,10^.

Although previous works have shown a difference in microbiome composition between hybrid and landrace maize varieties ^11^, the roles of seed-borne bacteria and their implications for plant development remain unclear. Some studies suggest that plant domestication may lead to a decrease in the transmission capability of the seed-borne microbiome ^12^, which would gradually reduce its composition and diversity of it in crop varieties used in intensive agriculture systems. Such a reduction in seed-borne microbial diversity may be linked to negative outcomes, like increased susceptibility to phytopathogenic diseases ^13^.

Metagenomic studies have also shown a strong correlation between reduced microbiome diversity and declines in both crop productivity and disease resistance ^14,15^. Within the plant core microbiome, several taxa play pivotal roles, such as *Pantoea*, renowned for their diverse plant-growth-promoting traits ^16,17^. Furthermore, specific genes in the bacteria, as it happens with Pseudomonads ^18^, can also enhance the capability of plant-microbe interactions ^19^. Notably, the maize microbiome exhibits a prevalence of families like Rhizobiaceae, Burkholderiaceae, and Microbacteriaceae, particularly during early plant development ^20^. These families are usually associated with traits such as phosphate solubilization and nitrogen metabolism ^21–24^.

With a rising emphasis on eco-friendly practices in sustainable agriculture, bioinoculants have been adopted over the last four decades, a market that grows at an annual rate of 10% worldwide ^25^. Hence, by combining the effects of environmentally compatible inoculants and microbiome knowledge that comes from omics analysis, we can alter the “stage 0” of the seed microbiome, focusing on increased crop productivity and protection ^3^ while also compensating for any potential reduction of seed-borne microbial diversity in intensive crop varieties. Currently, the most effective way to increase crop production is through the widespread use of chemical pesticides and fertilizers. However, this demand for chemical inputs increases dependency on imported products that compromise the sovereignty of several countries for food, fiber, and energy production ^26,27^. Furthermore, the intensive use of agrichemicals is also associated with groundwater contamination ^28^, human diseases ^29^, and death of pollinating insects ^30^. In this scenario, the adoption of sustainable practices based on plant growth-promoting bacteria (PGPB) constitutes an attractive alternative ^31^. PGPB typically promote plant growth through diverse mechanisms of action such as biofertilization, bioprotection, or biostimulation ^32^.

In the present work, we investigate the seed-borne bacteriomes of maize varieties derived from two distinct breeding strategies. We evaluated the microbiome of a commercial hybrid and a landrace maize variety, the latter originating from organic production in an agroforestry ecosystem. We analyzed the structure and diversity of these microbiomes, with particular emphasis on their differences and potential to promote plant growth.

## Methods

### Maize varieties

We used the “SHS 5050” double hybrid commercial variety (SH) and the “Sol da Manhã” homozygous landrace variety (SOL). SH is a hybrid variety obtained from a local Santa Helena retailer with production in northwest Minas Gerais - Brazil. This company works to improve maize yield and uniformity, dealing with a large-scale production area irrigated under a central pivot. The SOL variety was donated by the family of the agribusiness farmer Jamil Bráz Corinto to Prof. Samuel Kamphorst (UNILA, Brazil). This variety originated from agricultural growth corridors in Santo Antônio do Rio Verde, Goiás, Brazil. SOL was initially cultivated in indigenous lands and has been cultivated in agroforestry systems for over 15 years, a period during which it has undergone meticulous participatory mass selection. The cultivation of SOL is a collaborative effort supported by EMBRAPA and with involvement from the Movimento Camponês Popular (MCP).

### Planting

We washed maize seeds from both varieties with sterile distilled water five times and soaked them for five hours to break dormancy and standardize germination. We carefully distributed the seeds into sterile 2 L glass jars filled with 700 g of Basaplant substrate sieved through a 2 mm mesh and previously autoclaved for four one-hour cycles. We irrigated the substrate with 250 mL of autoclaved distilled water and divided it into three distinct sectors, each containing 10 seeds. We sealed the glass jars with cotton to allow gas exchange while maintaining the seedlings under axenic conditions throughout the incubation period. We conducted the experiment in a BOD incubator (100 µmol/m^2^/s), maintaining the temperature at 28°C “day” and 25°C “night”, following a 12-hour light/12-hour dark cycle for 7 days. We chose to work with germinated seeds so that the microbiome would mimic the real scenario of plant development in the soil instead of studying only the quiescent seed microbiome ^16,33^.

### 16S and metagenomic sequencing

We collected the rhizospheric soil by brushing it from the seedlings and conducted DNA extraction using the DNeasy PowerSoil Pro Kit (Qiagen) following the manufacturer’s instructions. We sequenced the microbiome DNA libraries using an Illumina NextSeq instrument at NGS (Piracicaba, Brazil). We also sequenced a DNA sample from the autoclaved substrate to serve as a negative control (“C1”) for contamination in shotgun sequencing. The raw sequencing reads were submitted to NCBI SRA under the BioProject PRJNA1069023. For 16S rRNA sequencing, we used the Wizard Genomic DNA Purification Kit (Promega) followed by sequencing using the Sanger method using the 27F (5′-AGAGTTTGATCCTGGCTCAG-3′) and 1492R (5′-TACGGYTACCTTGTTACGACTT-3′) universal primers ^34^.

We used QIIME2 v. 2023.5.1 ^35^ to process 16S metataxonomic data and remove data from mitochondrial and chloroplast 16S rRNA. We used the SILVA database Release 138 ^36^ to infer taxonomic classification.

For metagenome analysis, we used Bowtie2 v. 2.3.4.3 ^37^ and Samtools v. 1.9 ^38^ to remove fragments of maize DNA. We then evaluated sequence quality using FastQC v. 0.12.1 (http://www.bioinformatics.babraham.ac.uk/projects/fastqc/) and trimmed low-quality fragments using Trimmomatic v. 0.39 ^39^. The taxonomic distribution of the reads was conducted with Kaiju v. 1.6.2 ^40^.

For maximizing plant growth promotion capacity screening in our metagenomes, we inferred the microbiome biotechnological potential using an in-house database of genes associated with different plant growth promotion traits (i.e. nitrogen fixation, phosphate solubilization, phytases, ACC deaminase, and auxin production) (Pedrosa-Silva, Henaut-Jacobs, Venancio, in preparation). We constructed this database by selecting genes widely known for their plant growth promotion capabilities. We included copies of each gene from every different genera that are available in UniProt. Predicted microbiome genes were screened using this database with Usearch ^41^ with a 60% identity threshold.

### Metagenome-Assembled Genomes (MAGs)

We used SPAdes v. 3.15.5 ^42^ to make the metagenomics assembly using the “*--meta*” parameter. Assembly quality was inferred using QUAST v. 5.0.2 ^43^ to help choose the best assembly parameters based on assembly metrics. We retained only contigs longer than 1750 base pairs. MAGs were binned with MetaBAT2 v. 2.2.15 ^44^. We used the GTDB toolkit Release 08-RS214 ^45^ to infer the taxonomy and BUSCO v. 5.4.2 ^46^ to assess the completeness and duplication levels of the MAGs. The redundancy between the MAGs was estimated with pyani v. 0.2.12 and their genes were predicted with Prokka v. 1.13 ^47^. We adopted the MIMAG methodology ^48^ and retained only non-redundant MAGs considered High-quality or Finished for submission to Genbank.

## Results and discussion

### Metagenomics sequencing depth satisfactorily captured sample diversity

We started the analysis with the C1 control sample, which had 1881 operational taxonomic units (OTUs) (Table S1, Table S2). Most of these OTUs are exclusive to C1, suggesting that they result from residual DNA from bacteria killed during sterilization. Conversely, the OTUs from SH and SOL are remarkably more abundant than those of C1, most likely because of living bacteria in these samples. These results support the observation that the DNA reads sequenced from the SH and SOL samples were indeed from the seed-borne microbiome, with virtually no soil DNA contamination.

To assess the coverage of our data, we estimated the taxa per sample (Figure 1) and computed the Good’s coverage index for each sample (Table 1). These analyses demonstrated that our sequencing data achieved robust taxonomic coverage, capturing 80% of the microbial diversity at 10 million reads in both varieties. Notably, the Good’s index is in line with the rarefaction curves, with only a slight variation where SH exhibits marginally lower Good’s indexes in comparison to SOL. SH samples also exhibited a more delayed plateau phase (Figure 1), which is consistent with their greater number of singletons (Table 1) that probably originated from low-abundance bacteria that are unlikely to significantly contribute to the community structure. Nevertheless, we kept these singletons for downstream analysis because of the overall low bacterial DNA amounts in our samples.

**Table 1:**
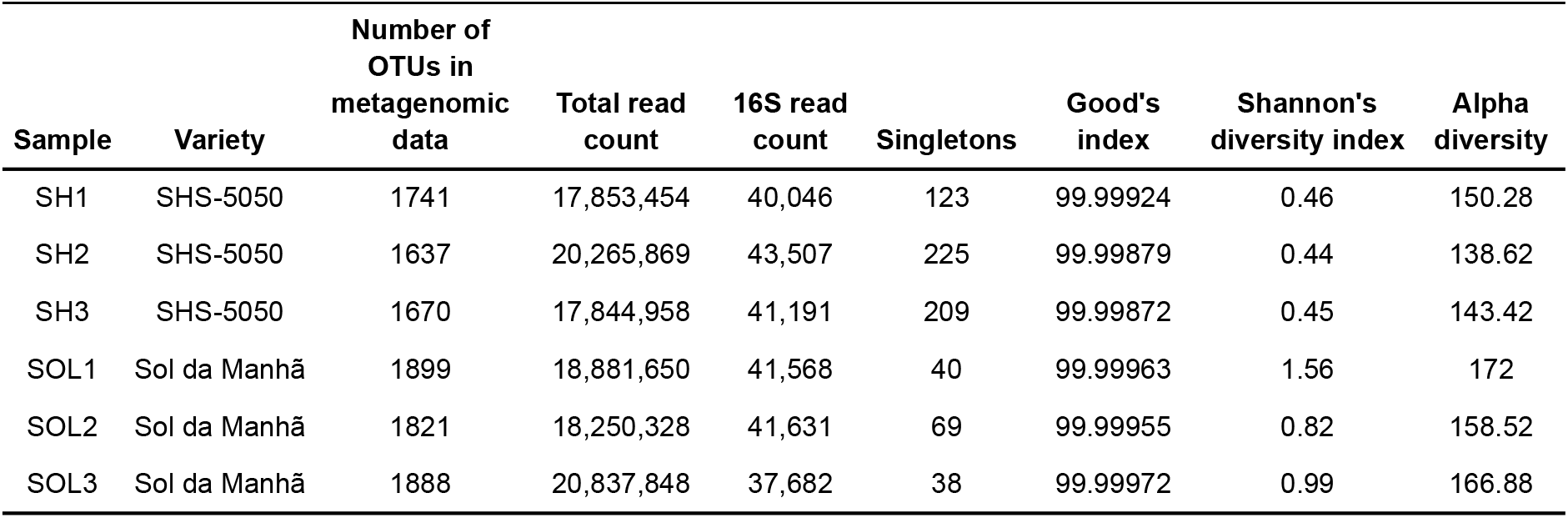
Complete sample information with sample identification based on the Methods description (Santa Helena variety = SH; Sol da Manhã variety = SOL).

**Figure 1:**
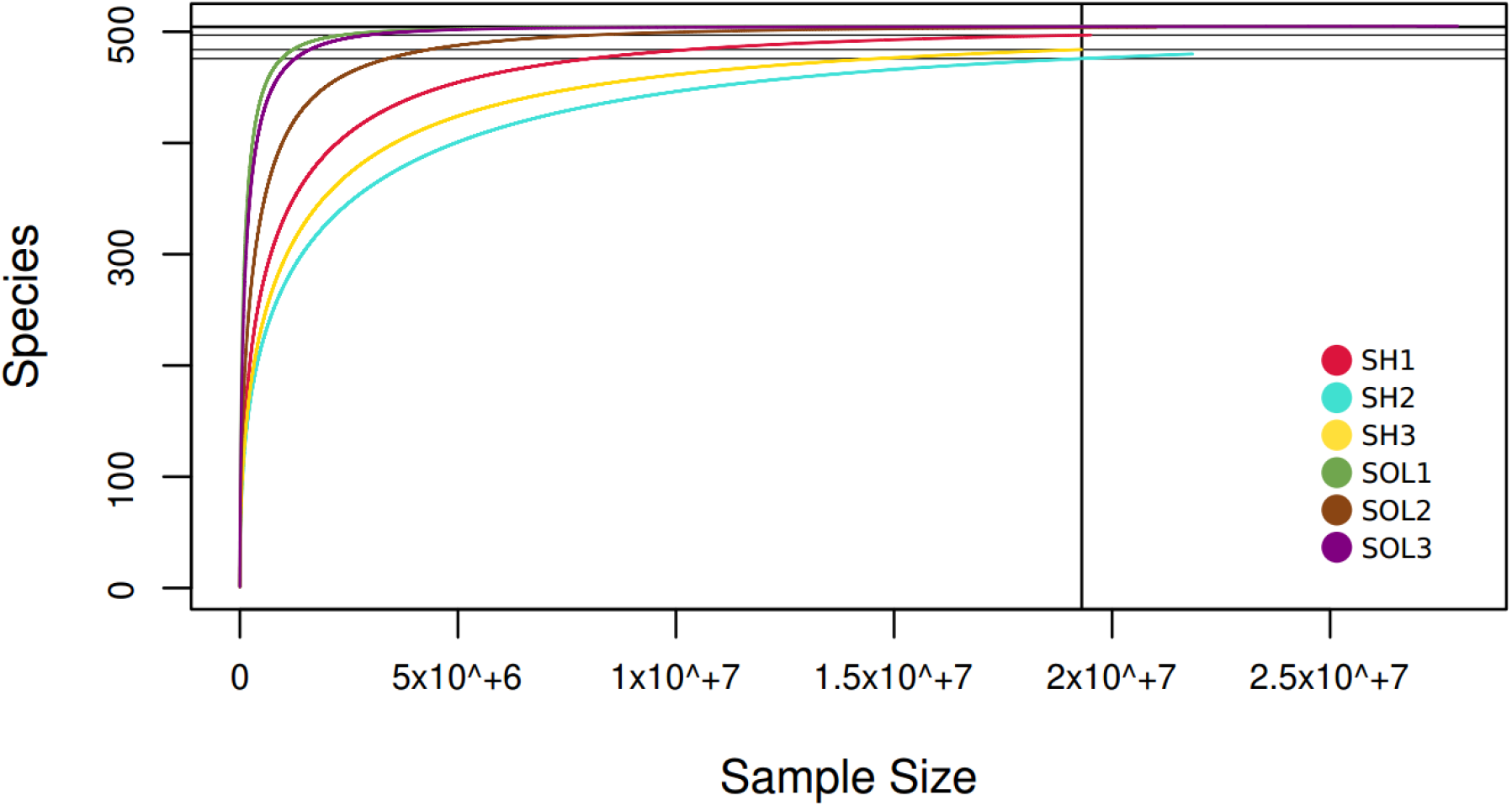
Saturation curve of OTUs found in each sample. Horizontal lines indicate the final number of species from each sample.

### The hybrid maize genotype harbors a homogeneous seed-borne bacteriome

We sequenced 124,744 and 120,881 16S rRNA partial gene sequences in SH and SOL samples, respectively (Table 1). In our 16S metataxonomic data, we observed a substantial abundance of the Burkholderiaceae family in SH and SOL, mainly from the *Burkholderia* and *Paraburkholderia* genera (Figure 2.B). Previous studies reported the prevalence of the Burkholderiaceae family in maize rhizosphere microbiomes ^11^. Importantly, *Burkholderia* species have been associated with nitrogen fixation and stress resistance in maize ^49,50^. One of the most frequent species in both maize varieties was *Burkholderia cenocepacia* (Table S1), which is part of the *Burkholderia cepacia* complex that has been widely reported as associated with plant growth promotion ^51^. The inoculation of *B. cenocepacia* improved maize production (cob length, number of grains per cob, grain weight, and 100-grain weight) in the field, with better results when combined with *Alcaligenes aquatilis* ^52^.

**Figure 2:**
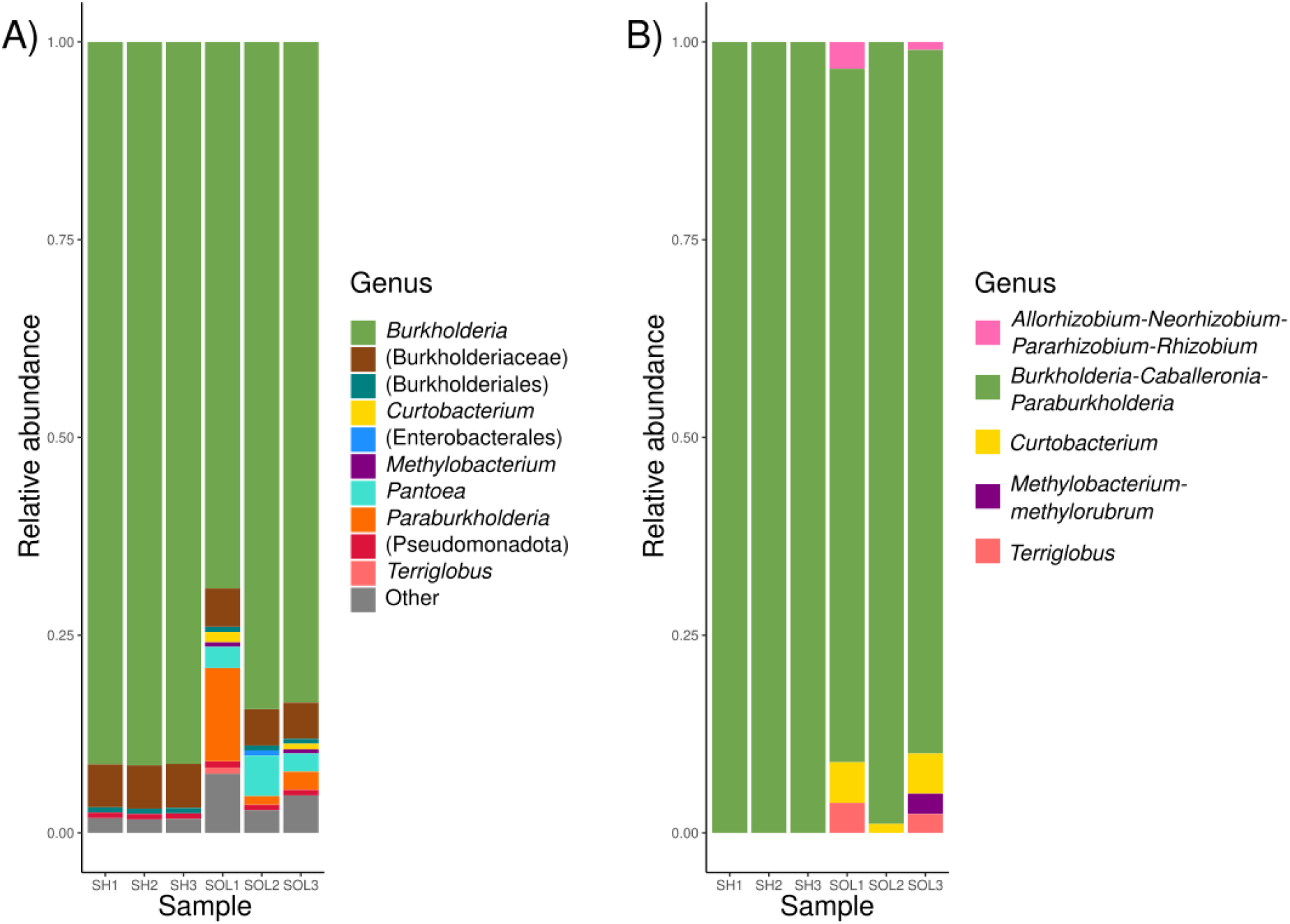
Relative abundance of OTUs. A) shotgun sequencing data (higher taxonomic levels were presented between parentheses), reads with less than 1% relative abundance across all samples, and reads without taxonomic assignment were collectively classified as “Other”; B) 16S rRNA data.

It is important to acknowledge that some *B. cenocepacia* strains are pathogenic to humans, particularly to immunocompromised individuals ^53^. Conversely, other *B. cenocepacia* strains are non-pathogenic and deemed safe ^54,55^. Hence, it is feasible to strategically incorporate non-pathogenic strains in inoculants or synthetic microbiomes to harness the beneficial features of *B. cenocepacia* as a PGPB, while mitigating potential risks to human health.

Burkholderiaceae accounted for over 90% of the SH microbiome. In addition, all classified reads from the SH 16S metataxonomic data were assigned to the *Burkholderia*-*Caballeronia*-*Paraburkholderia* genera (Figure 2b). In contrast, SOL samples displayed a substantial presence of various taxa and variable proportions of Burkholderiaceae in their composition. A Principal Component Analysis (PCA) further highlighted this distinction, with SH samples almost completely overlapping, while SOL samples were more dispersed (Figure 3). We can also see a separation of SOL1 from the other samples, which reflects the lower prevalence of *Burkholderia* in this specific sample.

**Figure 3:**
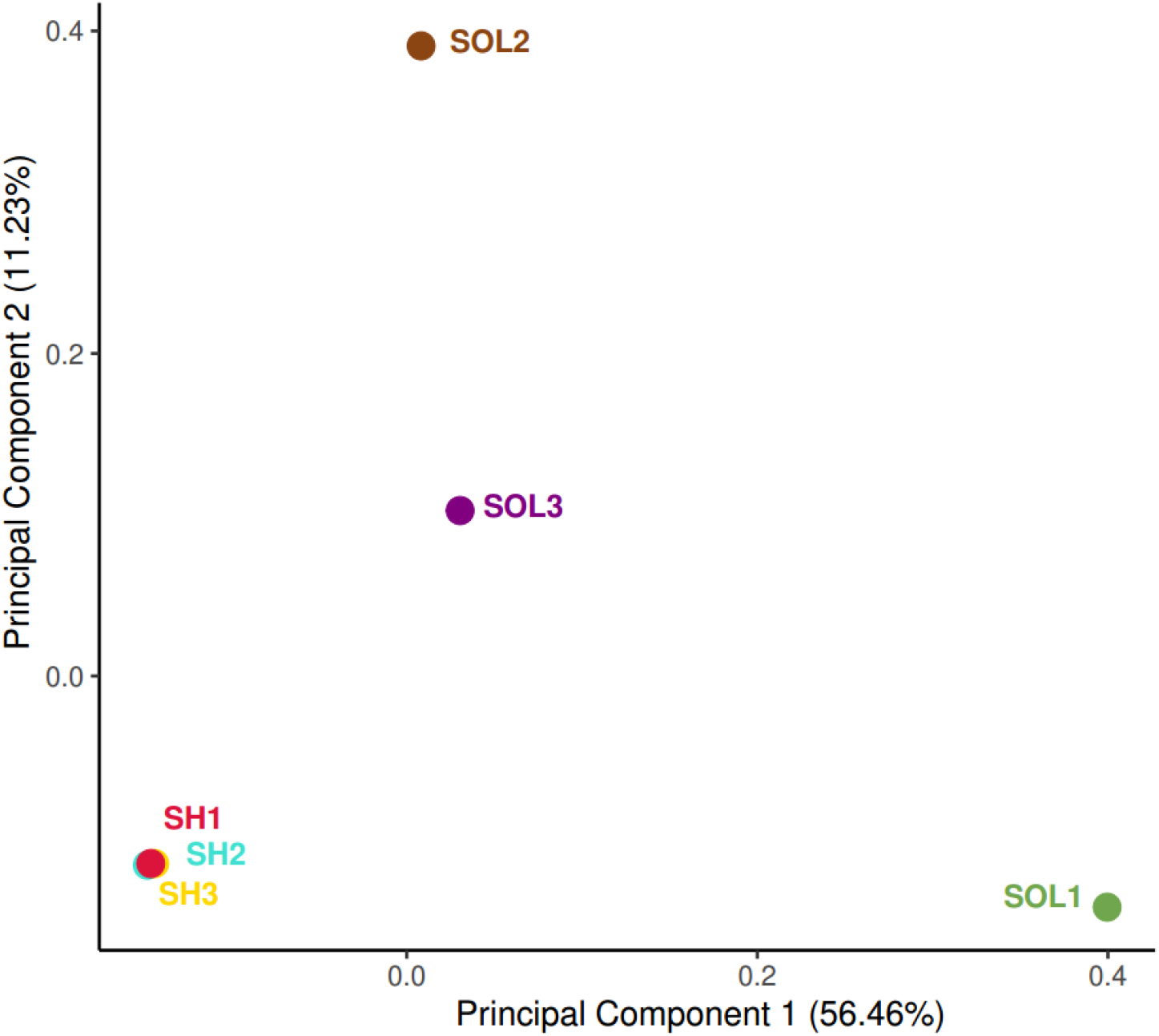
Principal Component Analysis performed with the OTU diversity and abundance across samples. The analysis was conducted with the VEGAN R package ^59^.

The homogeneity found in SH samples mirrors the inherent genetic uniformity of the hybrid variety. It is well-established that the plant genotype profoundly influences the microbiome composition and that intensive agriculture can reduce microbiome diversity ^56^. Prior research has demonstrated differences in microbiome recruitment in inbred maize varieties over time, with more recently developed germplasms recruiting fewer microbial taxa with the genetic capability for nitrogen fixation and larger populations of microorganisms that contribute to nitrogen loss ^57^. It is reasonable to assume that SH experienced a loss of microbiome diversity due to its long-term use in intensive agriculture when compared to SOL, which is also reflected in the high occurrence of singletons in those samples. Another critical aspect to consider is that the intensive use of chemical fertilizers and pesticides reduces the selective pressure on recruiting a supportive microbiome by plants ^58^, which might also be the case in commercial varieties like SH.

### The seed-borne microbiome harbors multiple phosphate solubilization genes but is likely unable to fix nitrogen

Bacteria can promote plant growth through various mechanisms, either directly or indirectly. Here we use a list of manually curated genes to investigate the presence of direct plant-growth promotion mechanisms, including nitrogen fixation, phosphate solubilization, ACC deaminase and auxin production.

We expected a greater abundance of nitrogen fixation genes in SOL because of the gene loss driven by extensive breeding and chemical nitrogen fertilization in varieties like SH ^57,60^. Contrary to our hypothesis, the *nifHDK* core nitrogenase genes were absent in both varieties. Nevertheless, we found *nifB* and *nifUS*, which are involved in the biosynthesis of the FeMo nitrogenase cofactor ^61,62^ and in providing the essential Fe-S clusters for the biosynthesis of FeMo-Co^63^, respectively. In SH samples, we found *nifQ*, responsible for molybdenum incorporation, which acts in conjunction with other genes (*nifB, nifNV*, and *nifE*) in the biosynthesis of the nitrogenase iron-molybdenum cofactor ^64,65^ (Figure 4). The relevance of these *nif* genes in the absence of the core nitrogenase subunits is yet to be clarified. However, it is important to emphasize that even non-intensive agricultural management involves nitrogen fertilization. This recurrent nitrogen supplementation could result in a high abundance of nitrogen in the soil, reducing selective pressure for retaining diazotrophic microorganisms in the seed-borne microbiome.

**Figure 4:**
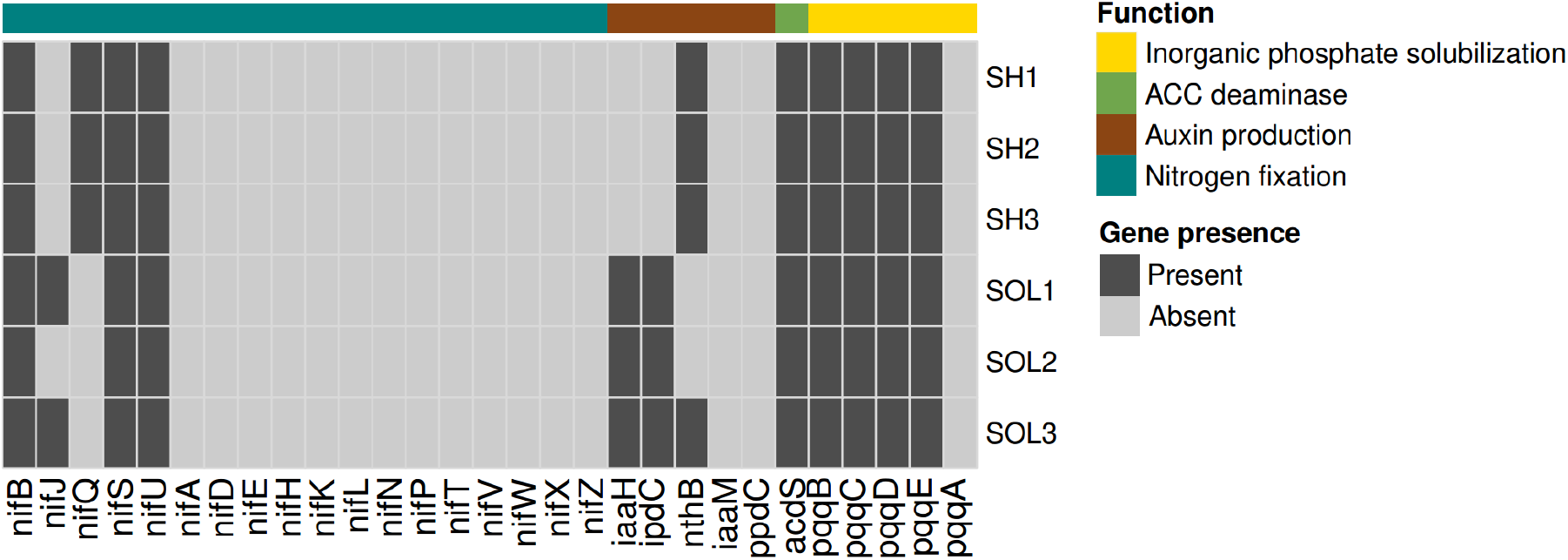
Functional heatmap of plant-growth promotion genes in each sample.

We found all the *pqq* genes for inorganic phosphate solubilization in all samples, except for the *pqqA* gene (not necessary for PQQ cofactor synthesis ^66^). Further, phytase genes are absent in metagenomes, supporting the dominance of inorganic phosphate solubilization in the seed-borne microbiome. The evolution of the seed-borne microbiome to prioritize the solubilization of inorganic over organic phosphate is compatible with the general availability of phosphorus in soils used in agriculture ^67^.

### Seed-borne microbiome may affect phytohormone production

We found distinctive patterns of auxin biosynthesis genes, with SH harboring the *nthB* gene, while SOL showed a dominance of *ipdC* and *iaaH*. These results show a much greater auxin biosynthesis potential in the SOL seed-borne microbiome. In this context, we can hypothesize that increased utilization of nitrogen fertilizers and intensive breeding can make auxin-producing microorganisms dispensable ^68,69^.

We also found the ACC deaminase gene *acdS* in all samples. ACC deaminases convert ACC in α-ketobutyrate and ammonia ^70,71^, lowering ethylene levels in the plant ^32^. Reduced ethylene levels can delay senescence ^72^ and improve plant resistance to stressful environments ^73^ and pathogen attacks ^74^. Hence, the presence of *acdS* could also be favored by selection in intensive agriculture systems where plants are often under reduced stress.

### Metagenome-assembled genomes (MAGs)

A total of 18 valid bacterial MAGs were recovered (Table 2). We identified ten high-quality MAGs that met the stringent criteria of at least 90% genome completeness and up to 5% contamination. There were five medium-quality draft MAGs, with 50 to 90% completeness and less than 10% contamination, and one low-quality draft MAG, with less than 50% genome completeness and more than 10% contamination. Two MAGs exhibited duplication levels exceeding 10% and were likely contaminated.

**Table 2:**
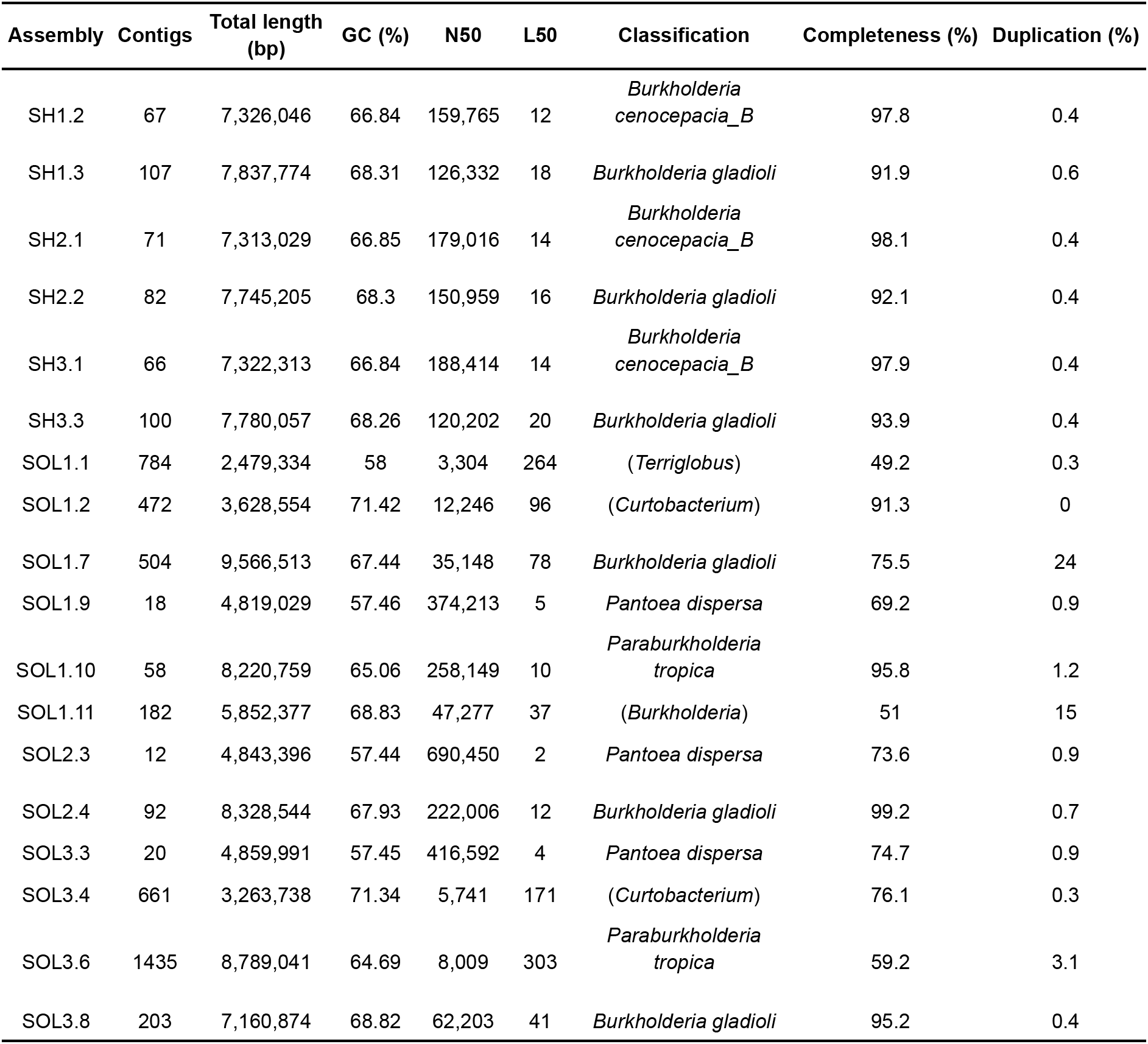
MAGs characteristics and metadata. Classification is only at the genus level and is presented between parentheses.

| Assembly | Contigs | Total length (bp) | GC (%) | N50 | L50 | Classification | Completeness (%) | Duplication (%) |
| --- | --- | --- | --- | --- | --- | --- | --- | --- |
| SH1.2 | 67 | 7,326,046 | 66.84 | 159,765 | 12 | *Burkholderia cenocepacia_B* | 97.8 | 0.4 |
| SH1.3 | 107 | 7,837,774 | 68.31 | 126,332 | 18 | *Burkholderia gladioli* | 91.9 | 0.6 |
| SH2.1 | 71 | 7,313,029 | 66.85 | 179,016 | 14 | *Burkholderia cenocepacia_B* | 98.1 | 0.4 |
| SH2.2 | 82 | 7,745,205 | 68.3 | 150,959 | 16 | *Burkholderia gladioli* | 92.1 | 0.4 |
| SH3.1 | 66 | 7,322,313 | 66.84 | 188,414 | 14 | *Burkholderia cenocepacia_B* | 97.9 | 0.4 |
| SH3.3 | 100 | 7,780,057 | 68.26 | 120,202 | 20 | *Burkholderia gladioli* | 93.9 | 0.4 |
| SOL1.1 | 784 | 2,479,334 | 58 | 3,304 | 264 | (*Terriglobus*) | 49.2 | 0.3 |
| SOL1.2 | 472 | 3,628,554 | 71.42 | 12,246 | 96 | (*Curtobacterium*) | 91.3 | 0 |
| SOL1.7 | 504 | 9,566,513 | 67.44 | 35,148 | 78 | *Burkholderia gladioli* | 75.5 | 24 |
| SOL1.9 | 18 | 4,819,029 | 57.46 | 374,213 | 5 | *Pantoea dispersa* | 69.2 | 0.9 |
| SOL1.10 | 58 | 8,220,759 | 65.06 | 258,149 | 10 | *Paraburkholderia tropica* | 95.8 | 1.2 |
| SOL1.11 | 182 | 5,852,377 | 68.83 | 47,277 | 37 | (*Burkholderia*) | 51 | 15 |
| SOL2.3 | 12 | 4,843,396 | 57.44 | 690,450 | 2 | *Pantoea dispersa* | 73.6 | 0.9 |
| SOL2.4 | 92 | 8,328,544 | 67.93 | 222,006 | 12 | *Burkholderia gladioli* | 99.2 | 0.7 |
| SOL3.3 | 20 | 4,859,991 | 57.45 | 416,592 | 4 | *Pantoea dispersa* | 74.7 | 0.9 |
| SOL3.4 | 661 | 3,263,738 | 71.34 | 5,741 | 171 | (*Curtobacterium*) | 76.1 | 0.3 |
| SOL3.6 | 1435 | 8,789,041 | 64.69 | 8,009 | 303 | *Paraburkholderia tropica* | 59.2 | 3.1 |
| SOL3.8 | 203 | 7,160,874 | 68.82 | 62,203 | 41 | *Burkholderia gladioli* | 95.2 | 0.4 |

Remarkably, we discovered four MAGs belonging to novel species, including two from the *Curtobacterium*, one from *Burkholderia*, and one from *Terriglobus*. These four novel MAGs were found in SOL samples, with SOL1.2 being classified as high-quality according to the MIMAG methodology ^48^. Furthermore, the two unknown *Curtobacterium* MAGs were 99% identical, supporting their affiliation to the same novel species.

It is reasonable to assume that identifying a high-quality MAG is more common for abundant strains. Thus, the independent recovery of the same MAG in different samples strongly supports its robust occurrence in the seed-borne microbiome. Species such as *Burkholderia gladioli*, found in all six samples, might have an essential role in the seed-borne microbiome. Importantly, given the ANI values greater than 99%, we probably found the same *B. gladioli* strain in all six samples. *B. gladioli* has been previously reported as a beneficial seed-borne bacteria, showing a good repertoire of phosphatases and no nitrogen fixation genes ^75^, which is coherent with the results presented here. *B. gladioli* has also been shown to enhance plant resistance to *Fusarium oxysporum* by inducing fungal cell death ^76^. We hypothesize that the presence of *B. gladioli* in all samples could represent a seed-borne biocontrol mechanism to counter phytopathogenic fungi during germination and early post-germination development.

Another interesting MAG is that of a *B. cenocepacia* strain found in all SH samples. *B. cenocepacia* is also a known biocontrol agent ^77^ generally associated with fusarium wilt protection. Furthermore, *B. cenocepacia* was already reported to be involved with *Fusarium* root rot protection by controlling other species of *Fusarium* that should work together with *F. oxysporum* ^*78*^. The co-occurrence of *B. cenocepacia* in the SH seed-borne microbiome with *B. gladioli* could be a signal of an evolving system of actively recruiting biocontrol agents by maize’s mother plant. In addition, our results also support the compatibility of these two strains to work together, either additively or synergistically. We believe that, with further validation, these strains could be used as part of a novel biocontrol strategy.

## Concluding remarks

In this study, we demonstrated that a landrace maize variety has a more diverse and less uniform (between samples) seed-borne microbiome than a hybrid commercial variety. We identified a higher phytohormone production potential in the landrace seed-borne microbiome. When exploring MAGs, we identified that some taxa are uniformly distributed between different varieties, supporting the existence of a core seed-borne microbiome. We also found an unexpected absence of *nif* genes in both varieties, which could be linked to the widespread use of nitrogen fertilization even in non-extensive agricultural environments. Understanding the dynamics of how seed-borne microbiomes evolve can guide the development of novel precision agriculture applications.

## Acknowledgments

This work was supported by Fundação Carlos Chagas Filho de Amparo à Pesquisa do Estado do Rio de Janeiro (FAPERJ; grants E-26/203.014/2018, E-26/201.142/2021, and E-26/201.117/2022), Coordenação de Aperfeiçoamento de Pessoal de Nível Superior - Brasil (CAPES; Finance Code 001), and Conselho Nacional de Desenvolvimento Científico e Tecnológico (CNPq). The funding agencies had no role in the design of the study and collection, analysis, and interpretation of data and in writing.

## Author contributions

Conceptualization: SH-J, FLO, TMV; Formal analysis: SH-J; Computational methodology: SH-J, FP-S; Wet lab methodology; BEBC; Writing – original draft: SH-J, TMV; Writing - review & editing: SH-J, BEBC, FLO, TMV; Funding acquisition: FLO, TMV; Project administration: FLO, TMV.

## Data availability

All the genomic sequences of this paper were submitted under the BioProject PRJNA1069023.

